# Public-good driven release of heterogeneous resources leads to genotypic diversification of an isogenic yeast population in melibiose

**DOI:** 10.1101/2021.04.12.439421

**Authors:** Anjali Mahilkar, Phaniendra Alugoju, Vijendra Kavatalkar, E. Rajeshkannan, Paike Jayadeva Bhat, Supreet Saini

## Abstract

Adaptive diversification of an isogenic population, and its molecular basis has been a subject of a number of studies in the last few years. Microbial populations offer a relatively convenient model system to study this question. In this context, an isogenic population of bacteria (*E. coli*, *B. subtilis*, and *Pseudomonas*) has been shown to lead to genetic diversification in the population, when propagated for a number of generations. This diversification is known to occur when the individuals in the population have access to two or more resources/environments, which are separated either temporally or spatially. Here, we report adaptive diversification in an isogenic population of yeast, *S. cerevisiae*, when propagated in an environment containing melibiose as the carbon source. The diversification is driven due to a public good, enzyme α-galactosidase, leading to hydrolysis of melibiose into two distinct resources, glucose and galactose. The diversification is driven by a mutations at a single locus, in the *GAL3* gene in the GAL/MEL regulon in the yeast.

## Introduction

Metabolic specialization can lead to diversification of an isogenic population. This phenomenon has been observed when diversification happens to (a) occupy the different niches available to a population [1, 2], (b) occupy new niches created by the population [3–6], or (c) occupy novel, previously unavailable, niches via evolution of metabolic innovation [7, 8]. In these cases, such specialization has been observed because of acquisition of a relatively small number of mutations [9–11]. The repeated observation of emergence of specialists and the relatively easy route in the sequence space facilitating this transition indicates that metabolic specialization is likely an important mode for creating genetic diversity.

Recently, it was shown that microbial populations occupy precise niches, when allowed to grow in spatially structured spaces [12]. In this context, the examples cited above demonstrate diversification when there is a spatial or temporal heterogeneity in how the resources are made available to the population. However, how diversification takes place in well-mixed environments containing multiple resources is not well understood.

Diversification can be geometrically viewed as an isogenic population starting from a single point in a valley of a landscape, and different parts of a population going up different peaks. Therefore, diversification should, in this view, becoming increasingly harder as populations move up an adaptive peak [13]. However, this view is one of static landscapes. Ecological interactions lead to dynamic evolution of the environment and the associated landscapes too. In this context, it was shown that evolution in an environment first leads to competitive adaptation, followed by diversification [14].

Coexistence of multiple genotypes in an environment can also be facilitated by metabolic trade [15, 16]. It has been demonstrated experimentally and theoretically that an auxotrophic pair of genotype/species, trading essential metabolites, can grow faster than the prototroph parent [15, 17]. However, how, starting from an isogenic prototrophic population, we can achieve a population split to take place is not well understood.

One particular manifestation of metabolic trade between two or more species is emergence of a cheater population [18]. In such a context, some members of an isogenic population pay a cost for production of a public resource. This leads to emergence of cheaters in the population, which do not contribute towards production of the public resource, but gain from benefits from the resource produced by co-operators in the population. Because the cheaters have a higher fitness than the co-operators, but yet cannot survive in the absence of co-operators – a stable coexistence results [19, 20]. Presence of cheaters in an environment has also been shown to have a positive effect in preserving biodiversity in a unstructured space competition experiments between bacteria [21]. This is because presence of cheaters decreases the fitness of the co-operators, thus, allowing other species to not be eliminated in the resulting environment. However, effects of cheaters on being able to influence the fitness of cooperators are likely to be dependent on the precise environment in which fitness is tested [22].

Impact of cooperation and cheating in dictating structures of population has also been investigated using game theory. These studies have demonstrated that if interactions between participating species/genotypes can be represented via Hawk-Dove, Snowdrift games, a stable coexistence results [23, 24]. This game-like representation was applied to *Pseudomonas* and *Klebsiella*, and coexistence shown on spatial and uniform environments [25].

In yeast, it has been shown that when feeding on a non-simple sugar, sucrose – the population structure influences the fate of the population (collapse or coexistence) [26, 27]. In this case, hydrolysis of the disaccharide leads to release of glucose and fructose. Interestingly, the hydrolysis takes places in the periplasm, leading to a small fraction (∼0.01) of the hydrolysed sugars to diffuse into the extracellular membrane and being available as a public resource. The dynamics of growth of the two participating genotypes (co-operator and cheater) in this case is dictated by the initial frequency of the participating genotypes in the population [28]. Growth on sucrose as a carbon source has two characteristic features. First, growth dynamics of yeast on glucose and fructose, the constituent monosaccharides of sucrose, are quite similar, particularly in non-fermenting conditions [29, 30]. Second, only a small fraction of the total resource (the two hydrolysed monosaccharides) is made available as a public resource.

Several *S. cerevisiae* strains can also grow on melibiose as a carbon source. Melibiose is a disaccharide of glucose and galactose, and is hydrolysed into its constituent monosaccharides by the action of an extracellular α-galactosidase enzyme encoded by the gene *MEL1* [31]. Of the two monosaccharides, *S. cerevisiae* has evolved the ability to consume glucose rapidly, and has evolved to have a galactose utilization regulon, which is extremely sensitive to the amounts of glucose in the environment [32]. Expression of genes necessary for galactose utilization is activated by the Gal4p transcription factor [33, 34]. In the absence of galactose, Gal4p is sequestered by the Gal80p, and promoters of the gal regulon are in the OFF state. The Gal80p-dependent repression is relieved in the presence of galactose, when the signal transducer Gal3p or Gal1p bind Gal80p, hence, freeing up Gal4p to activate gene expression from promoters of the gal regulon. Gal1p, in addition to its role as a signal transducer, also acts as a kinase, necessary for galactose utilization in the cell [35, 36] (**Figure 1**).

**Figure 1.**
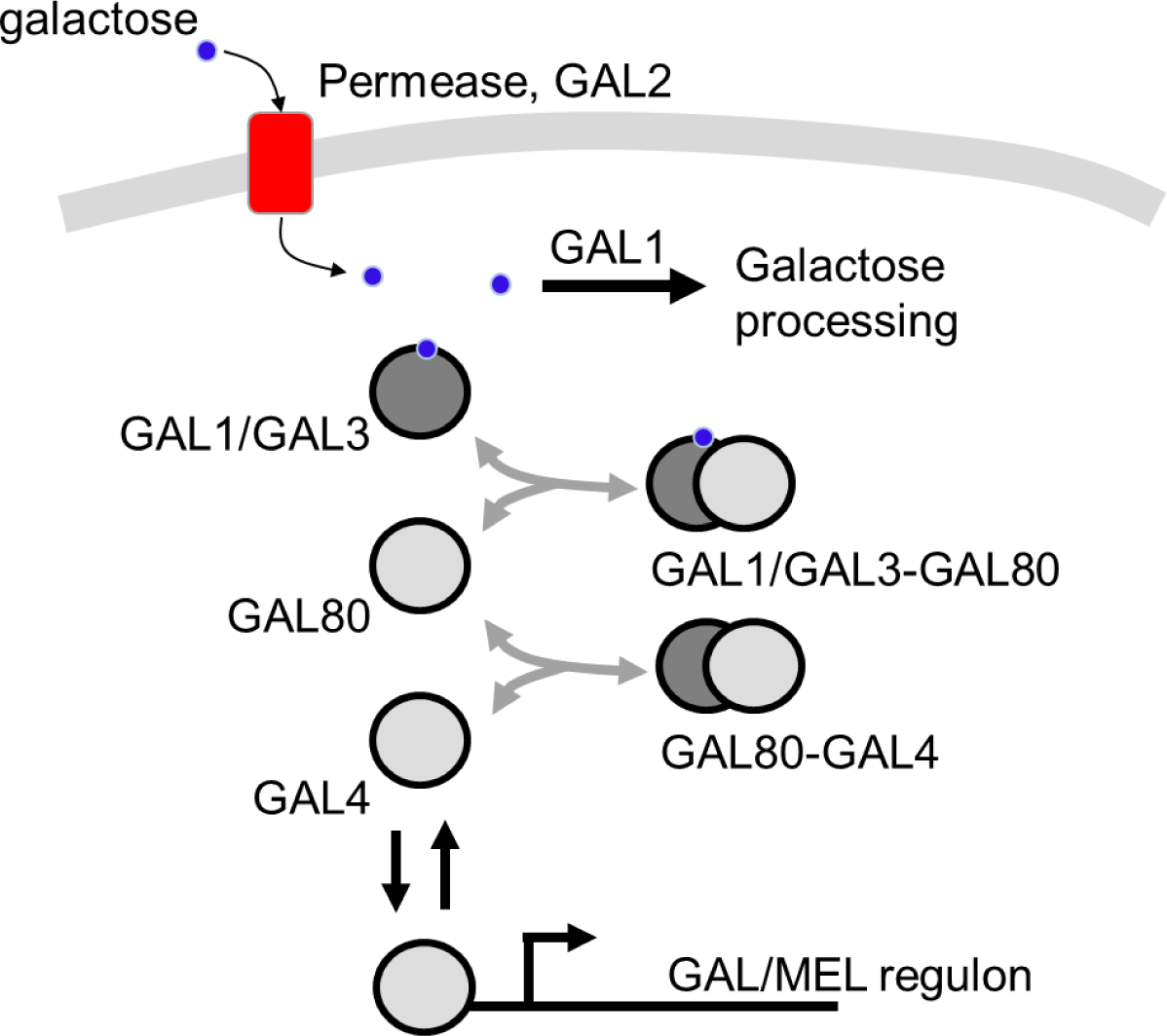
Gal/Mel network. Genes in the GAL/MEL regulon are activated by the transcription factor Gal4p. Gal4p is sequestered by the protein Gal80p, which binds to Gal4p to form the Gal80-Gal4 protein complex. Gal80p-dependent repression is relieved when, in the presence of galactose, Gal3p/Gal1p bind Gal80p, thus freeing Gal4p for activation of the network. Galactose is imported into the cell via the permease Gal2p. Intracellular galactose is processed via metabolic enzymes Gal1p, Gal7p, and Gal10p.

This system is different from the sucrose utilization system in two different aspects. First, the disaccharide is hydrolysed in the extracellular environment. Hence, all of glucose and galactose so produced is a public resource. Second, kinetics of growth (duration of lag phase and growth rate) of *S. cerevisiae* on glucose and galactose are quite different from each other (**Figure 2**).

**Figure 2.**
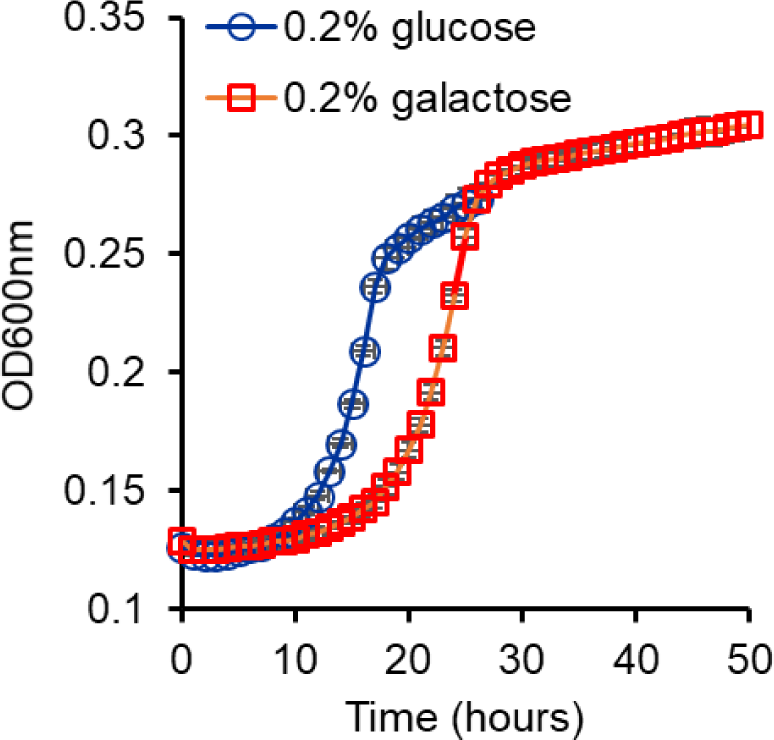
Growth kinetics of *S. cerevisiae* in glucose and galactose. Growth rate on glucose is 0.0438 h^-1^ and that in galactose is 0.0406 h^-1^. The duration of lag in glucose is 11 h and that in galactose is 17 h. Growth rate and the duration of lag were determined by taking log of the OD600 values and plotting against time. The slope of the linear part of the trajectory calculated as the growth rate of the culture. The intercept of the linear part in this curve with the log of OD600 at t = 0 was used as a measure of the duration of the lag phase.

In this work, we use the melibiose utilization system in *S. cerevisiae* as a model system in which processing of a resource (melibiose) by a public good (Mel1p) can lead to the population splitting into two distinct genotypes. We demonstrate that during growth on melibiose, metabolic heterogeneity is observed in an isogenic population during the exponential phase of growth. By serial sub-culturing to maintain the cells in the state of heterogeneity, growth for a few hundred generations leads to the population genetically splitting into two distinct phenotypes. The two phenotypes/genotypes are distinguished by the colony size on solid melibiose media, and also by growth dynamics in glucose and galactose. Sequencing results show that the coexistence is maintained via polymorphism at the *GAL3* locus. Overall, these results show that simple genetic changes can facilitate diversification of an isogenic population into two distinct genotypes even in spatially unstructured environments. This diversification is driven by dynamic release of a public good (Mel1p) which leads to release of heterogeneous carbon sources (glucose and galactose).

## Materials and Methods

### Strain used

A SC644 diploid strain (*MATa MEL1ade1 ile trp1-HIII ura3-52*) was used in this study. BY4743 was used for the competition experiment. BY4742 ΔGAL3::KanMX4 (Euroscraf) was used for complementation experiments.

### Growth kinetics

Glycerol-lactate pre-grown strains were plated onto SCM agar plates (containing sugar concentration as defined). The plates were thereafter incubated at 30°C for 3-4 days. From the plates, colonies were randomly selected from the agar plates and subjected to two rounds of serial passage in appropriate media (as described in the results section). The resulting cultures were then washed with SCM and then growth curves were initiated with an initial optical density of 0.1 in SCM containing sugar(s) at an appropriate concentration as described. Three replicates of culture were transferred to a 96-well plate and OD was measured for every one hour until the cultures reach stationary phase. The plates were overlaid with a *Breathe Easy* membranes (Sigma) to prevent evaporation. To calculate the growth rate, log(OD), in the exponential phase of growth, was plotted against time. The slope of the straight line fit was calculated as the growth rate of the strain. The x-axis value (time) where this straight line intercepts y = log(initial OD) was taken to be the duration of the lag phase of growth.

### Sugar estimation

Cell free liquid samples were analysed for melibiose and galactose by high performance liquid chromatography (HPLC) using JASCO (PU-2080) system and Biorad-aminex HPX-87H column. Analytes were detected using JASCO refractive index detector (RI-2031plus) by keeping polarity +16. Standard solutions of galactose and melibiose were prepared using HPLC grade water from a concentration range of 0.1 mg/ml to 10 mg/ml. The standard graph was prepared using the following conditions: mobile phase 10 mM H_2_SO_4_, injection volume 10 μl, temperature 65 deg C, and a flow rate of 0.5 ml/min for 20 min.

Glucose was estimated in the extracellular media by analysing the cell-free liquid samples using the GOD-POD kit (Atlas Medical). 10 μl of the sample or standard was mixed with 1000 μl of glucose mono-reagent and incubated at 37 deg C for a duration of 30 min. The intensity of the red dye was quantified spectrophotometrically at a wavelength of 507 nm. Standard graph of glucose was prepared using HPLC grade water from a concentration range of 0.1 mg/ml to 10 mg/ml.

### Sporulation

Cells freshly grown on YPD plates were patched on about 1 cm^2^ area to a freshly prepared GNA pre-sporulation plate for 1 day at 30°C. After growth for 24 hours, the cells from the GNA pre-sporulation plate were re-patched to another freshly prepared GNA pre-sporulation plate for 24 h at 30°C. After 24 h of growth, the cells from the GNA pre-sporulation plate were re-patched to a sporulation medium plate, and incubated at 25 deg C for 5 days followed by incubation at 30 deg C for three days [37].

### Experiments with 2-deoxy galactose (2DG)

Yeast ancestral and strains evolved 1% melibiose were inoculated into Gly/lac medium and incubate for 48-72 hrs. These Gly/lac pregrown cultures were inoculated into 5 mL of CSM containing 1% melibiose and incubate at 30 deg C. Cells were harvested (from respective ancestral and evolved cultures) from exponential phase, serially diluted with PBS and thereafter, plated onto different concentrations of 2-Dexoygalactose (2DG) (0%, 0.3 μM, and 0.6 μM) containing Gly/lac plates along with Gly/lac plates. Plates were incubated at 30 deg C for 3-4 days. The number of colonies that grew on gly/lac and 2-Dexoygalactose were counted and the percentage of Gal-positive cells calculated for both ancestral and evolved strains.

### Evolution experiment

Yeast diploid SC644 strain was used for the evolution experiment. A single colony was grown in liquid CSM galactose medium (inducing medium) for 24 h at 30°C on a rotary shaker at 250 rpm. In order to initiate evolving cultures 50 μL of this overnight grown pre-culture was inoculated at a dilution of 1:100 into fresh selection medium (CSM containing 1% melibiose). Three parallel populations lines were evolved in high sugar medium i.e., CSM containing 1% melibiose. Populations were propagated in 5 mL of liquid medium (30°C, 250 rpm) within 25 x 150 mm borosilicate tubes. After every 24 h of growth, (i.e., after an average of 6.64 generations/day) 50 μL of each culture was transferred to 5 mL of fresh medium containing respective concentrations of melibiose daily for up to 400 generations. Every 100 generations, aliquots of each evolving population were suspended in 30% v/v glycerol and store at –80°C for further use.

### Colony size analysis

Plate Images were taken in *UVITECH* gel documentation unit in white light at an exposure of 800 ms, 3x zoom and 900 focus using the *Essential V6* software across days starting from Day 2 of plating to Day 10. The colony size was measured using CellProfiler 3.1.8 using the automated pipeline described in [38]. All colonies between pixels 6 to 95 were considered during the analysis.

### Sequencing

Sequencing of the promoter regions and the coding sequences was done using the following primers. For *GAL1*, primers 5’ – TTA ACT GCT CAT TGC TAT AT – 3’ and 5’ – AAA AGA AGT ATA CTT ATA AT - 3’ were used; for *GAL3*, primers 5’ – GCT TTT ACT ATT ATC TTC TA – 3’ and 5’ – TTG TTC GTA CAA ACA AGT AC – 3’ were used; for *GAL4*, primers 5’ – GGA CCC TGA CGG CGA CAC AG – 3’ and 5’ – CAT TTT ACT CTT TTT TTG GG – 3’ were used; for *GAL80*, primers 5’ – CAG ATG GAA TCC CTT CCA TA – 3’ and 5’ – GCA CTG GGG GCC AAG CAC AG – 3’ were used; and for *MEL1*, primers 5’ – GTC GAC TTC TAA GTA AAC AC – 3’ and 5’ – TGC TTT GCT CAA CAA TAA GC – 3’ were used. *MEL1* sequence was taken from a previous report [39].

The diploid evolved strain was plated on melibiose plates to isolate small and large colonies. Three large and three small colonies were picked up from a melibiose plates and the diploid sporulated. The four isolated haploids were isolated from each of the six colonies and grown on YPD plates. The colonies of the haploids were then grown in liquid YPF plates for 6-8 hours at 30 deg C. The cells were harvested, and their genomic DNA isolated. The DNA sequences were amplified by PCR. Sequencing was done by Eurofins Scientific.

### GAL3 cloning

The mutant *GAL3* gene was amplified using the primers pSC034 (CGA GTC GAA TTC AAT ACA AAC GTT CCA ATA) and pSC038 (AAG CTT GAG TAA ACT TTT AAT ATT TAA) from the large colony of evolved E1 line from the melibiose plate and cloned into the plasmid pYJM [40] between the EcoRI and HindIII cut-sites. The ancestral allele was amplified from the ancestor using the same primer set and cloned as described above. The resulting plasmids pYJM-GAL3* and pYJM-GAL3 and pYJM were transformed into BY4742 ΔGAL3::KanMX4 (Euroscraf) strain.

### Competition experiment

BY4743 and the ancestral strain were grown separately in non-inducing non-repressing conditions till saturation. Roughly 10^6^ cells of each genotype were transferred to a tube containing 1% melibiose, and allowed to grow for 24 hours. The culture (at t = 0 at the beginning of the experiment; and at t = 24 hours) was plated on YPD and ura-trp-double dropout plates to quantify the relative frequency of the two genotypes. The relative fitness was calculated using the formula below, as described in [41],

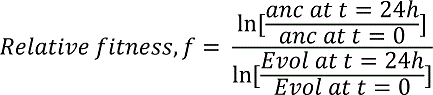

Where, anc. at t = 24h refers to the CFU count of the ancestral strain at time 24 h. The relative fitness of the evolved strain with respect to the strain BY4743 was calculated similarly.

### α-galactosidase enzyme activity

Extracellular α-galactosidase assay was performed to determine the expression level of the Mel1p, as described previously [42]. Yeast strains were grown in synthetic complete medium containing gly/lac up to saturation. The cultures were then sub-cultured in synthetic complete media containing 1% galactose to an initial OD of 0.05. The cultures were then allowed to grow till an OD of 1.00. A volume of 1ml of each culture was centrifuged and extracellular α-galactosidase activity of the supernatant was determined as follows. 120µl of the supernatant was mixed with 360 μl of assay buffer (2 volumes of 0.5 m sodium acetate, pH 4.5, and 1 volume of 100 mm p-nitrophenyl α-D-galactopyranoside (Sigma)). The reaction was incubated at 30°C for 5 h and terminated by adding 520 μl of stop buffer (1 M sodium carbonate). Enzyme amounts were then determined by measuring the absorbance at 410 nm. Triplicate samples were taken for the analysis and results represent average of at least three independent experiments with standard deviation.

### Cost-benefit modelling

In the generalist strategy, the cells can acquire mutations that eliminate glucose-dependent repression of galactose genes. In such a scenario, each individual cell uses glucose and galactose simultaneously, while exporting Mel1p. Since all cells behave metabolically identically in this strategy, we call this a generalist strategy. Each cell consumes equal amounts of glucose and galactose, which is equal to the amount of melibiose broken down by it. We start with modelling the benefit gained by a cell, in terms of fitness, as a function of sugar concentration.

At steady state in a chemostat, let the rate of melibiose hydrolysis per time by Mel1p released per cell be *k.* Therefore, the amount of glucose and galactose produced by hydrolysis to be consumed per time is also *k* each. Let us assume that the benefit conferred to the cell upon metabolising one molecule of glucose (or galactose) is *b*.

We assume that the benefit conferred to a cell upon consumption of *k* molecules of a substrate is of the form,

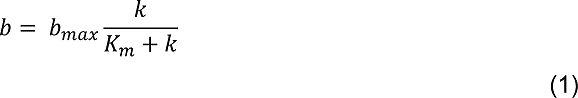

Where, *bmax* is the maximum possible benefit conferred as *k* approaches values much greater than half-maximum substrate amounts *Km.* In both the strategies we place the constraint that the maximum cellular flux that can be processed in a cell is *kmax*.

Since in the generalist case, the total number of molecules of the substrate consumed per time is *2k,* the benefit conferred to the cell can be written as,

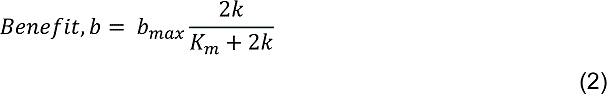

On the other hand, utilization of carbon substrate requires investment in the form of synthesis of appropriate enzymes and commitment of cellular resources (like, ribosomes, amino acids) towards enzyme synthesis. This cost is proportional to the number of substrate molecules being utilized. Moreover, suppose each enzyme molecule in the pathway processes *a* substrate molecules per time, then, the total cost can be represented as,

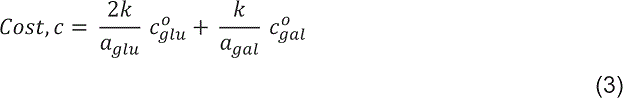

Where, *a_glu_* is the number of molecules processed per enzyme molecule in the glucose-utilization pathway, and *agal* is the number of substrate molecules per enzyme molecule in the galactose to glucose-6-phophate pathway. Note that from galactose-6-pathway, the processing of carbon is via a common metabolic path, and hence, *2k* molecules are added in the cost term.

The fitness of the cell is therefore given as,

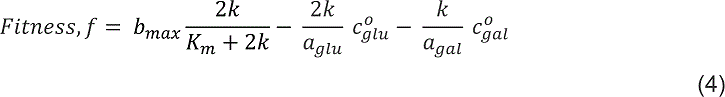

Alternatively, the population may evolve such that different cells adopt different strategies, with the goal that the fitness of the population is maximized. Such a strategy, where two or more different metabolic states optimize fitness, is referred to as a specialist strategy.

In this strategy, one fraction of the population hydrolysis melibiose into glucose and galactose. A fraction of the population is a cheater population, which does not contribute towards *MEL1* production, but instead utilizes glucose released from melibiose hydrolysis. As a result, the *MEL1-*producing fraction of the population consumes galactose for growth, and keep producing *MEL1* for continued hydrolysis of melibiose. The cheaters are preserved in the population because of their higher fitness, and the galactose users (ancestral cells) are maintained since they are essential for melibiose hydrolysis. Such a coexistence of two distinct genotypes has been observed in microbial system.

In a population of size *N,* let the fraction of co-operators (ancestral cells) by *x.* Therefore, the fraction of cheaters is (1-*x*). Let rate of hydrolysis by *k*, due to *MEL1* released by each co-operator cell. Therefore, total breakdown of melibiose per time = (*Nx*)*k*, and consequently, the amount of glucose and galactose released per time also equals (*Nx*)*k*.

Assuming that the galactose users consume galactose, the amount of galactose per cell is *k*. Similarly, assuming that the cheaters consume glucose, the amount of glucose per cell is *^kx^/1-x*. Therefore, as defined earlier, the fitness of co-operators is,

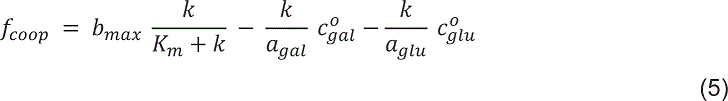

Similarly, the fitness of the cheaters cells can be quantified as,

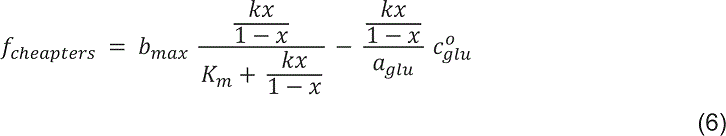

Now, for coexistence of co-operators and cheaters, the two should have equal fitness. All the parameters described above are inherent cellular properties, except for the hydrolysis rate *k*. It is a function of (1) the production rate of α-galactosidase, *po*, (2) The degradation rate of α-galactosidase, *kd*, and (c) processing rate of by α-galactosidase per enzyme, which is dependent on the concentration of melibiose, *α*.

The enzyme dynamics in the extracellular environment can be described as,

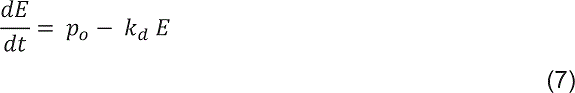

Where, *E* is the enzyme concentration in the environment. At steady state *E = ^po^/_kd_*. Thefore, the hydrolysis rate, *k = Eα*.

## Results

### Dynamics of growth in melibiose

Melibiose is a disaccharide of glucose and galactose. The enzyme Mel1p, which splits melibiose into the constituent monosaccharides is induced by GAL4p transcription factor, in the presence of galactose. Presence of glucose in the media represses expression of the *GAL* regulon, as well as that of *MEL1* [43]. Whether splitting of melibiose leads to sufficient glucose to successfully repress expression from the galactose regulon is not known. Logically, this situation is identical to that of lactose utilization in *E. coli*, with the only difference being that, in *E. coli*, the disaccharide is split into the monosaccharides inside the cell [44].

Growth on melibiose takes place after an uncharacteristically long lag phase. The dynamics of growth on increasing concentrations of melibiose is as shown in **Figure 3A**. The lag duration and the rate of growth are tunable parameters in this growth curve, and decrease and increase respectively, with increasing melibiose concentration. As compared with growth on equal amounts of carbon, the lag durations associated with glucose and galactose are much smaller compared to that of melibiose. It is not known whether this lag is due to slow induction of the cells (as they transition from a *MEL* OFF to a *MEL* ON state), or because of the glucose-induced repression?

**Figure 3.**
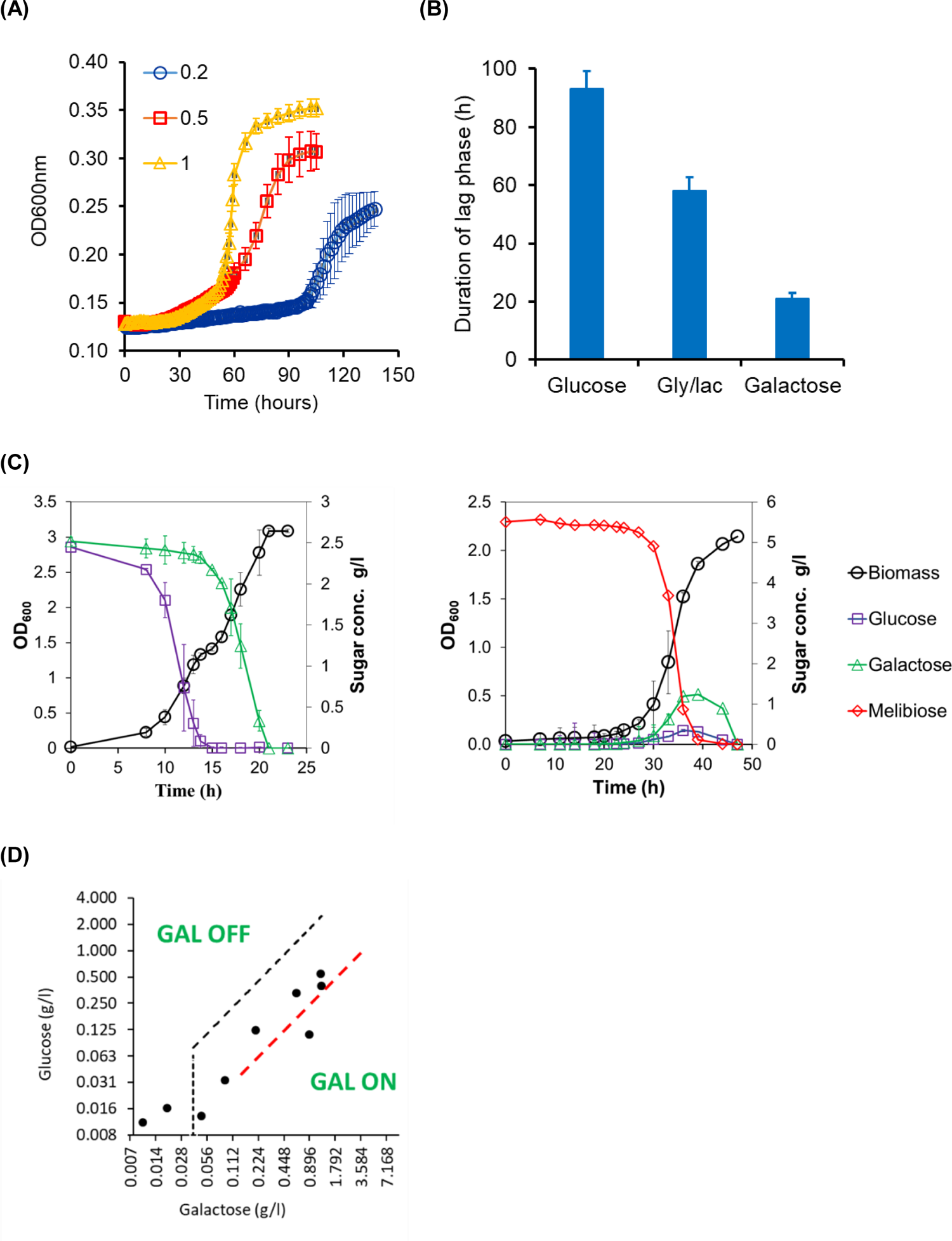
**(A)** Growth kinetics of *S. cerevisiae* in different melibiose concentrations. Cells were grown in non-inducing non-repressing conditions (gly/lac) and then sub-cultured in melibiose-containing media. Average of three runs and the standard deviation is reported. The duration of the lag phase was 42 h, 58 h, and 98 h for cultures grown at 1, 0.5, and 0.2 % melibiose respectively. The corresponding values of the growth rate were found to be 0.0385 h^-1^, 0.0145 h^-1^, and 0.01 h^-1^. **(B)** Duration of the lag phase depends on the pre-inoculum conditions. Lag phase duration in melibiose is longest when the cells are brought from glucose and shortest when the cells are introduced into melibiose media from galactose containing media. **(C)** Growth and sugar profiles when cells are grown on glucose-galactose mixture (left) and melibiose (right). Growth on melibiose is characterized by a long delay and simultaneous utilization of the two sugars. As a result, there is no daiuxy when cells are growing on melibiose. The transition from glucose to galactose in a glucose-galactose mixture (left) is clearly indicated by the sugar profiles in the media, and also by the growth dynamics. **(D)** The glucose-galactose concentrations in the extracellular media, during growth on melibiose, lie in a window of concentration range where cellular metabolic state is heterogeneous. That is, a part of the population has GAL genes in the ON state, and a part of the population has GAL genes in the OFF state. The dotted curves (black and red) indicate the decision front, either side of which the state of the cells is homogeneous (i.e. all cells have GAL genes OFF on the left of the decision front; and all cells have GAL genes in the ON state to the right of the decision front).

The duration of the lag is also a function of the initial state of the system. When cells from (a) an OFF state (in glucose), (b) an ON state (in galactose), and (c) neutral state (glycerol-lactate) with respect to GAL gene induction are introduced in media containing melibiose, the duration of the lag is strongly dependent on the environmental history of the cells. Cells transitioned from galactose exhibited the shortest lag, while those from glucose exhibited the longest lag phase, among the three conditions (**Figure 3B**).

Growth on melibiose is qualitatively different from that on glucose-galactose mixture. As shown in **Figure 3C**, when grown on a glucose-galactose mixture, the cells first utilize glucose and then transition to galactose. This transition is characterized by the classic diauxy phase. However, when cells are grown on melibiose, there is an uncharacteristic long lag phase, which is followed by co-utilization of glucose and galactose. In the log phase of growth, glucose and galactose accumulate in the extracellular media. The accumulation of the two sugars is unequal quantitatively, and the relative amounts of the two sugars lie in the window of concentration where cellular metabolic commitment, at a population level, is not unimodal (**Figure 3D**) [45].

### Cells growing in melibiose exhibit metabolic heterogeneity

Due to the design of the regulatory network dictating melibiose utilization, the metabolic strategy at a single-cell resolution, which maximizes growth rate of the population during lag phase, is not apparent. Hydrolysis of melibiose leads to glucose and galactose being released in the extracellular environment. One possible strategy can be that, between the two monosaccharides, an individual cell preferentially utilizes glucose first. As a result, the genes responsible for galactose utilization (including *MEL1*) are repressed. Once the glucose from the media is exhausted, the cell transitions to utilizing galactose. However, utilization of galactose triggers expression of *MEL1* resulting in release of glucose in the environment. Thus, the cell again switches back to the preferred carbon source glucose. However, such a strategy involves periodic shifting from one carbon source to another. Due to asynchrony in the population, at a single-cell resolution, however, at any given instant, there is metabolic heterogeneity. However, regular transitioning from one carbon source to another not only requires continuous adjustment in gene expression patterns, but also transitioning from one state to another at regular intervals. Both these factors have been demonstrated to contribute towards fitness costs, in terms of causing a reduction in microbial growth rates [46–48].

As against this, the other possibility of growth in melibiose suggests that during log phase of growth, the population splits into two distinct states. One fraction of the population utilizes galactose, and thus releases Mel1p in the environment for hydrolysis. In the remaining fraction of the population, the galactose network is in the OFF state, and cells grow by utilizing the glucose released as a result of hydrolysis. In such a scenario, at any instant in the population, an individual cell is present in one of the two metabolic states. Such a strategy, where an isogenic population splits into two phenotypic groups, has been observed in both bacteria [49] and yeast [50], when placed in environments with more than one carbon source.

The heterogeneity in the metabolic state of a population growing in melibiose can be tested by adding 2-deoxy-galactose (2DG), a non-metabolizable analog of galactose, to a cellular population. Upon addition, all cells which express Gal1p convert 2DG to a toxic intermediate metabolite and are thereafter killed [51]. Only cells which have the GAL system (Gal1p) in the OFF state survive addition of 2DG to the media. At the level of a single cell, the population exhibits a distribution of levels of GAL1 expression. Addition of a particular concentration of 2DG (0.3 μM) to the growth media places a threshold in terms of the cellular amounts of Gal1p. Any cell with Gal1p amount greater than this threshold is killed off, in presence of 2DG. If the concentration of 2DG in the media is increased, the threshold Gal1p amount decreases (**Figure 4A**). Thus, by studying the survival rates of a population at different 2DG concentrations, we can estimate the Gal1p distribution in that population.

**Figure 4.**
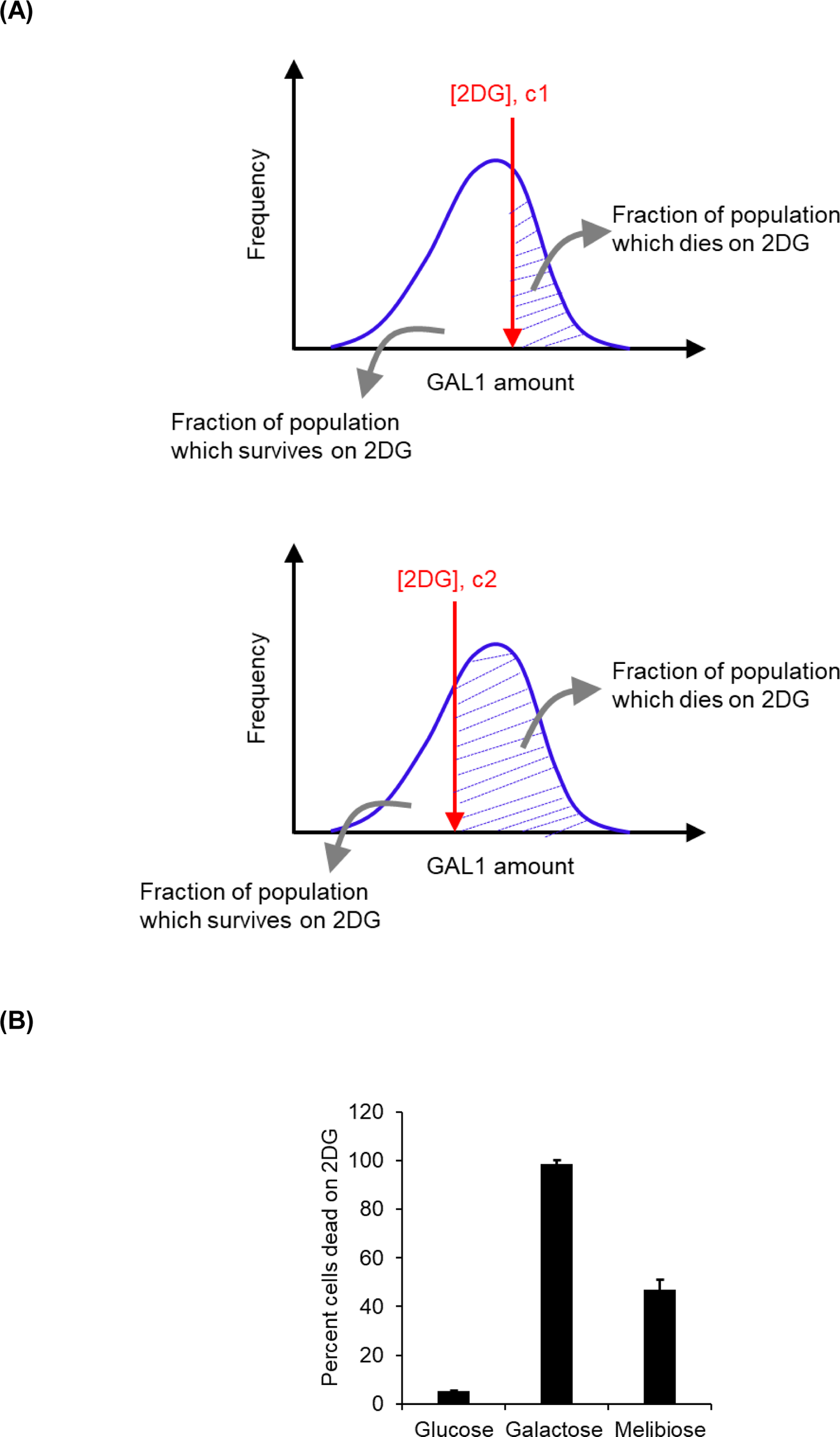
**(A)** Imagine a population of *S. cerevisiae* with a distribution of intracellular GAL1 levels. When spread on YPD plates with c1 concentration of 2DG, a GAL1 threshold is placed on the population. No cell with GAL1 level above the concentration do not survive on this plate. When the identical population is spread on a plate with a higher concentration of 2DG (c2), the GAL1 threshold is lowered. **(B) *S. cerevisiae* growing on 1% melibiose exhibits metabolic heterogeneity.** This heterogeneity is not exhibited when the population is growing on 1% glucose or 1% galactose. Wild type cells were grown to saturation in gly/lac media, and sub-cultured in respective media to an OD600 of 0.005. Cells were allowed to grow to an of 0.2 and plated on plates containing 2DG. Controls were spread on YPD plates. The experiment was performed in triplicate and the average and standard deviation are reported.

Adding 2DG to cells growing in glucose does not compromise cellular survival (>98% of the cells survive). However, addition of 2DG to cells in the mid-log phase of growth in galactose leads to nearly 100% cells being killed (**Figure 4**). When cells from the mid-log phase of growth in melibiose are taken and plated on a media containing 2DG, only around 50% of the cells survive. This is compared to almost 100% survival in the lag phase, and almost zero percent survival in the stationary phase of growth, in melibiose. These results demonstrate that during log phase of growth in melibiose, a population demonstrates metabolic heterogeneity.

In addition to the metabolic heterogeneity observed in the liquid media, when plated on melibiose, the population exhibits heterogeneity in the colony size distribution on melibiose. As shown in the **Figure 5**, the colony size distribution on glucose or galactose plates can be described using a normal distribution. However, the colony size distribution, on plates containing melibiose, is bimodal.

**Figure 5.**
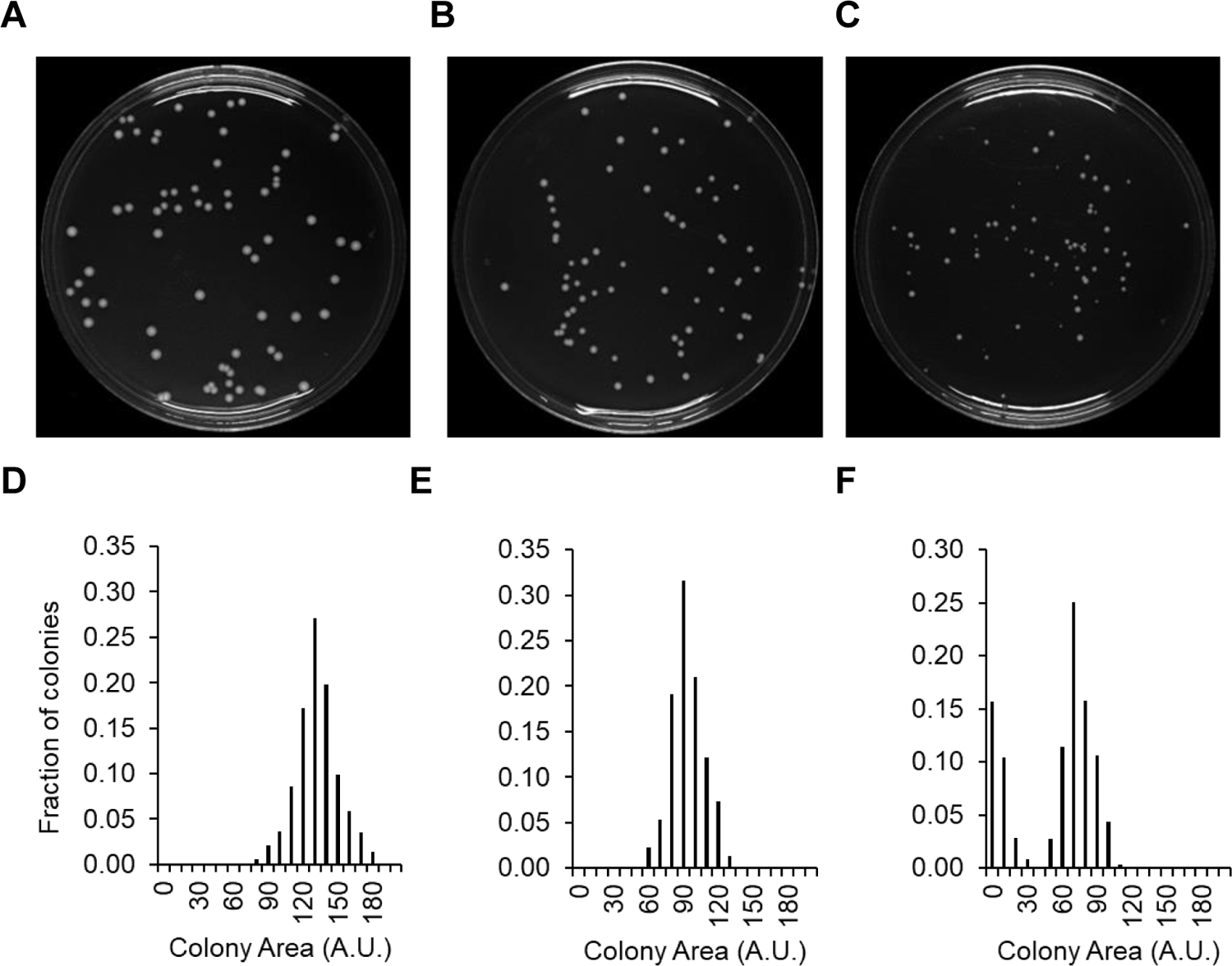
Colony size heterogeneity on melibiose plates. Cells grown in gly/lac were plated on plates containing glucose (A), galactose (B), and melibiose (C) as the carbon source. Each sugar was present at a concentration of 1%, and colony size distribution after 60 h of growth is shown. The colony size data was quantified and the distributions are shown for glucose (D), galactose (E), and melibiose (F). Distributions on glucose and galactose plates are represented by a normal distribution. Distribution on melibiose is bimodal.

Cells from a large and a small colony were transferred to and grown in gly/lac to saturation. The cells were then transferred to melibiose to a starting OD600 of 0.005 and the kinetics of growth monitored. In such a scenario, the two populations exhibit identical growth kinetics (**Figure 6**). This demonstrates that the heterogeneity in the colony size on melibiose is phenotypic in nature. The differences in colony size and their lack of correlation with the available resources (area on the plate) are likely due to the wide variation in the time at which an individual cell switches ON the melibiose utilization system. As a result, when cells growing in galactose are plated on melibiose, the heterogeneity in the colony size distribution is lost (**Figure 7**).

**Figure 6.**
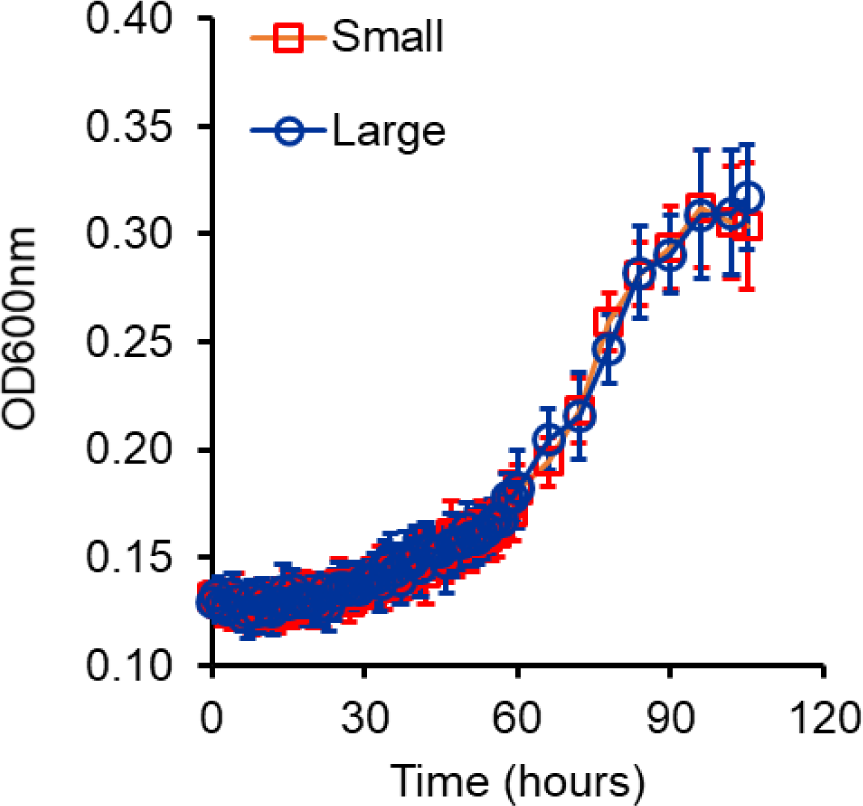
Small and large colonies on melibiose plates exhibit identical growth kinetics, once propagated through gly/lac. Cells from large and small colonies were suspended in gly/lac media and grown to saturation. Cells from the saturated culture were then sub-cultured in 1% melibiose media to an initial OD600 of 0.005. Growth kinetics of the cultures was then monitored at 30 deg C. Growth kinetics of six small and six large colonies was analysed. The average and standard deviation is presented in the figure above.

**Figure 7.**
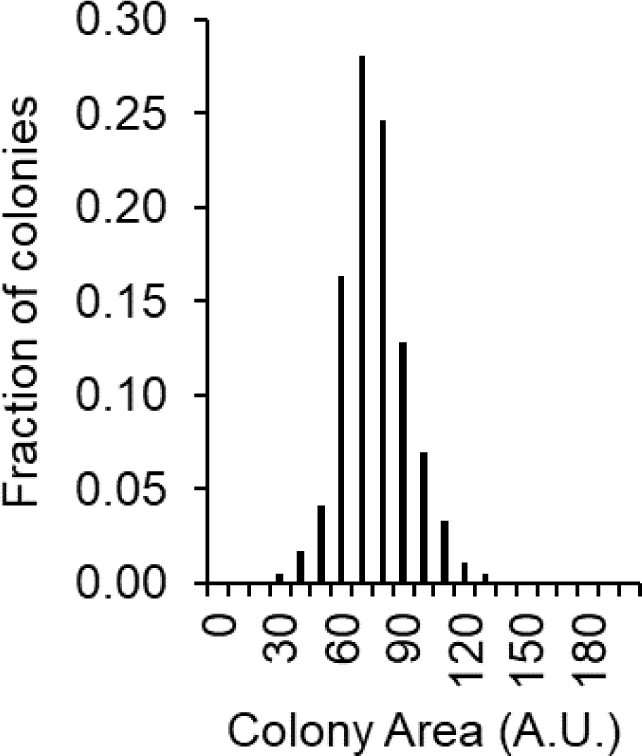
Colony size heterogeneity is not observed when cells are plated on melibiose from a galactose environment. Wild type cells were grown in 0.5% galactose to saturation and thereafter transferred to 1% melibiose plates for single colony. After 60 hours of growth, size of >500 colonies was measured and the frequency distribution of the size plotted.

### Cost-benefit model predicts that population diversification can be an adaptive strategy in high melibiose environments

Since there is metabolic heterogeneity in the wild-type population, when grown in melibiose, we ask the following question. If evolved in melibiose for a few hundred generations, does the metabolic heterogeneity observed collapse or get exaggerated? The two possible outcomes can both be argued as follows.

First, evolution in melibiose can be expected to lead to collapse in the heterogeneity by acquisition of mutation(s) which permit the cell to co-utilize glucose and galactose together. If such a generalist has a greater fitness than specialist populations, the population will not diversify genetically. Collapse of hierarchy of sugar utilization is known to take place even when microorganisms are evolved in precise sugar environments for a prolonged duration [52].

On the other hand, on evolution in melibiose environment, the phenotypic heterogeneity could evolve into a genetic heterogeneity. In such a scenario, the glucose users in the population would evolve to become better adapted to utilize glucose, whereas the galactose users will evolve to become galactose specialists. Such a genetic split will permit the two genotypes to coexist in the population.

To test the possibility of the two adaptive solutions, we develop a phenomenological mathematical model based on cost-benefit analysis which optimizes the growth rate of the culture under the two adaptive solutions. The logic of the model is follows. In the generalist adaptive solution, mutation(s) permit co-utilization of glucose and galactose. To derive benefit from the two sugars, the cell pays a cost to synthesize the necessary enzymes. However, an individual cell cannot process more than a specific amount of carbon flux per unit time. Such constraints are imposed by cellular physiology [53].

On the other hand, in the specialist adaptive solution, we hypothesize that a fraction of the population (*x*) utilizes galactose for growth and splits melibiose. The remaining fraction (1-*x*) grows on glucose produced as a result of this hydrolysis. Each cell type pays cost and derives benefit in accordance with the carbon source it is utilizing for growth. Since the two metabolic strategies coexist in the solution, during the log phase of growth, the model is solved for the value(s) of *x*, for which the two fractions have an equal growth rate/fitness.

We then compare the growth rates facilitated by each of the two strategies in the melibiose environment. As shown in the **Figure 8**, the best growth rate is facilitated by the generalist strategy when the parameter, *p*, (*^k_cat_s^/k_d_*) is small. The parameter *p* comprises of *kcat*, which is the enzyme catalytic activity for *MEL1*, *s* is the concentration of substrate melibiose present in the environment, and *kd* is the degradation rate of the *MEL1* protein in the extracellular environment. Thus, overall, the parameter *p* can be seen to be a proxy of the amount of hydrolysis taking place in the extracellular environment per unit time. After a critical value of the resource availability, the specialist strategy confers higher fitness to the population growing in melibiose.

**Figure 8.**
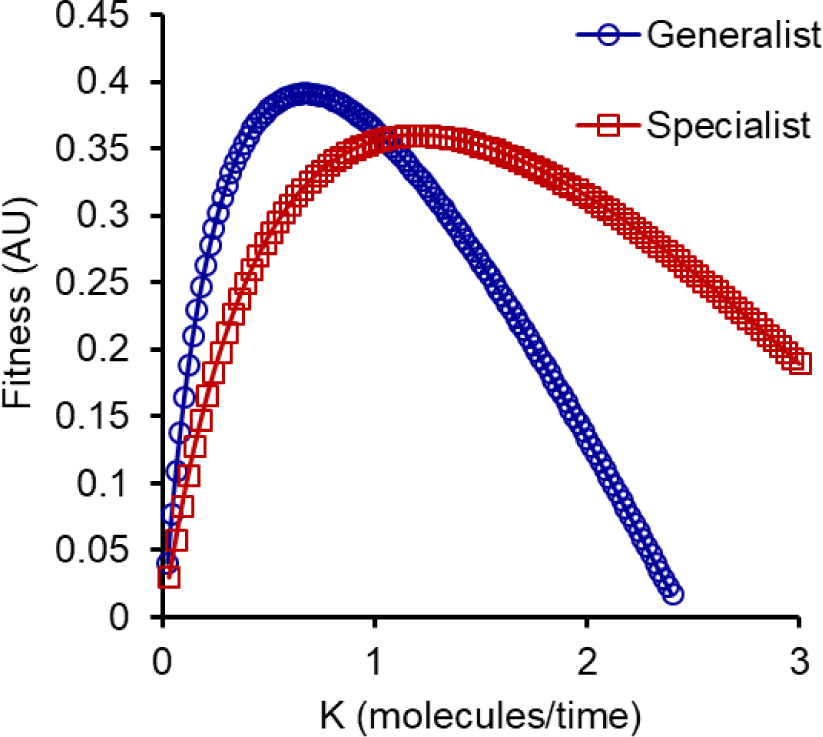
Cost benefit model predicts that specialists in glucose- and galactose utilization can evolve in high melibiose environments. The x-axis, *K*, equals (*^k_cat_s^/k_d_*) and represents the relative rate of melibiose hydrolysis in the surrounding environments.

This predicts that at high rates of melibiose hydrolysis, splitting the population into two specialist groups will lead to a higher fitness compared to a generalist strategy. The converse is true when the rate of melibiose hydrolysis is low. We note that in the context of melibiose utilization, the parameters *kd* and *kcat* have not been characterized experimentally, and their values are unknown. In this context, the ‘high’ and ‘low’ concentrations of glucose at which specialists and generalist, respectively, emerge as the optimal strategy are relative in nature.

Although the model does not comment on the access, in the sequence space, to the two solutions (generalist and specialist), we assume that the two solutions can be reached by an evolving population. How a population of isogenic individuals can adaptively evolve into a group of co-dependent fractions remains an open question [17].

When we pose the specialist groups in the population, we assume that one group exclusively uses glucose and the other uses galactose only. Intermediate strategies are also possible as an adaptive strategy, where one group exhibits a higher propensity for glucose, and the other for galactose. Any such intermediate strategy yields similar results, at different qualitative values.

### Evolution in high melibiose environments

We evolve yeast populations in melibiose environment for a duration of 400 generations in an environment containing two percent melibiose. Three independent lines were evolved as part of the experiment. The evolution experiment was carried so that cells growing in mid-log phase were transferred from the culture tube to a tube with fresh media. The transfer was thus done at a time in the growth phase when the population exhibited maximum metabolic heterogeneity.

After evolution for 400 generations in media containing 1% melibiose, we compare the growth curves for the ancestral and compare that with the evolved lines. The phenotype tested of all three evolved lines are indistinguishable from each other. As expected, the evolved lines exhibit a short lag phase and a higher growth rate as compared to the ancestor strain (**Figure 9**).

**Figure 9.**
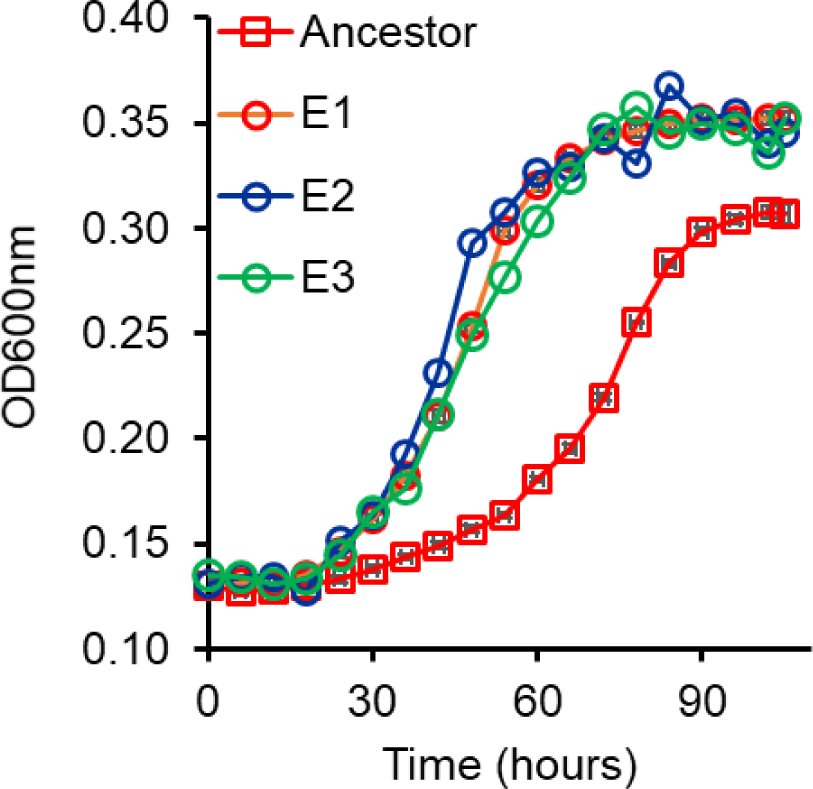
Kinetic of growth in melibiose of the three evolved lines. Cells were grown in gly/lac to saturation and sub-cultured in 1% melibiose to an initial OD600 of 0.005. Kinetics of growth was monitored at 30 deg C every 6 hours. Line 1 was taken forward for further analysis.

The beneficial mutations acquired during the evolutionary experiment help the cells exhibit a faster growth in glucose and galactose individually, too. That is, the evolved lines demonstrate a faster growth in glucose and galactose, as compared to the ancestor (**Figure 10**). This is itself is not a surprising result. Evolution in defined carbon environments with a single carbon source has been demonstrated to have little antagonistic effects [54]. In fact, evolution in one carbon environment has been reported to lead to pleiotropic benefits in other carbon environments too [55]. On the other hand, trade-offs between a number of aspects of physiology are well characterized too [56]. In the context of this experiment, since melibiose is comprised of glucose and galactose, it is perhaps not surprising that we do not observe any antagonism, in the form of compromised growth on glucose or galactose, in the strains evolved in melibiose. From the three lines evolved in this experiment, line E1 was further analysed in this work.

**Figure 10.**
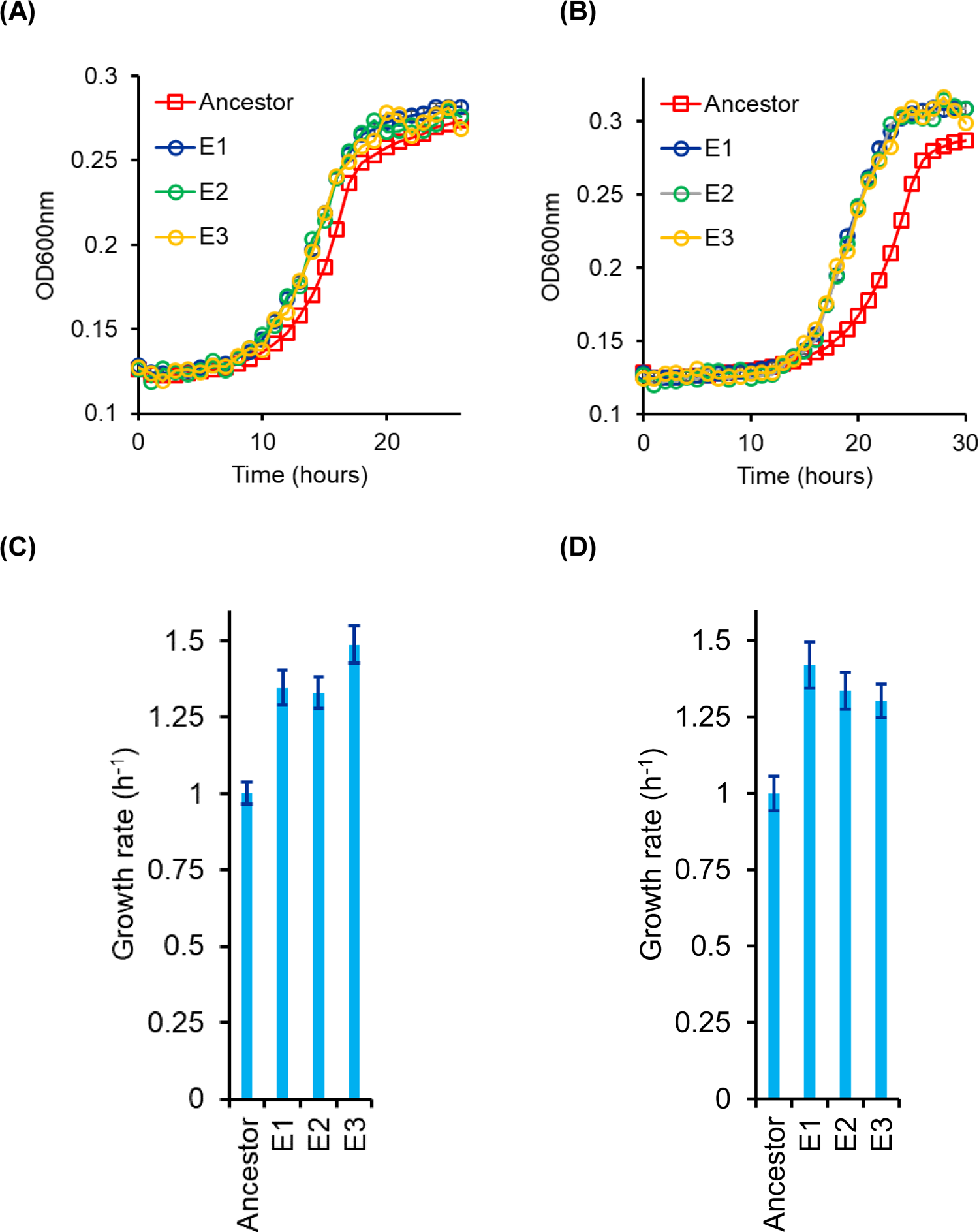
Growth kinetics of evolved lines in glucose (A) and galactose (B). Lines E1, E2, and E3 are the three independently evolved lines. (C) and (D) represent the growth rates in the exponential phase for the three lines for growth in 1% glucose and 1% galactose respectively. The growth rates of the evolved lines are normalized with respect to the ancestor’s growth rate in glucose (C) and galactose (D).

### Evolution for 400 generations in 1% melibiose leads to exaggerated metabolic heterogeneity

Gal1p amounts at a single-cell resolution exhibit a distribution, which can be quantified by adding different concentrations of 2DG to the media, and studying cell survival. We add a high concentration of 2DG (0.6 μM) to a culture of the ancestral cells in the mid-log phase of growth (OD 2) in 0.5% melibiose. At this concentration, only ∼3% cells survive. We call these cells ‘glucose specialists’, since they express minimal levels of Gal1p. At the same concentration of 2DG, however, approximately 8% of the evolved cells were able to survive. Thus, after evolution for 400 generations, the ‘glucose specialists’ in the population increased by almost three-fold.

In order to detect galactose specialists, to ancestral population growing in the mid-log phase in melibiose, we add a low concentration of 2DG (0.3 μM). This concentration only kills cells with fully induced *GAL* system (when grown in 1% galactose). In the ancestral population, this concentration killed approximately 2% of the population. This fraction of the population was estimated by subtracting the number of CFU that survived on 2DG from the total CFUs on plates without 2DG. We call this fraction of the population as ‘galactose specialists’. In the evolved lines, however, ∼9.5% of the population was killed at this low concentration of 2DG, indicating that the fraction of the population in the metabolic state of ‘galactose specialists’ had increased (**Figure 11**).

**Figure 11.**
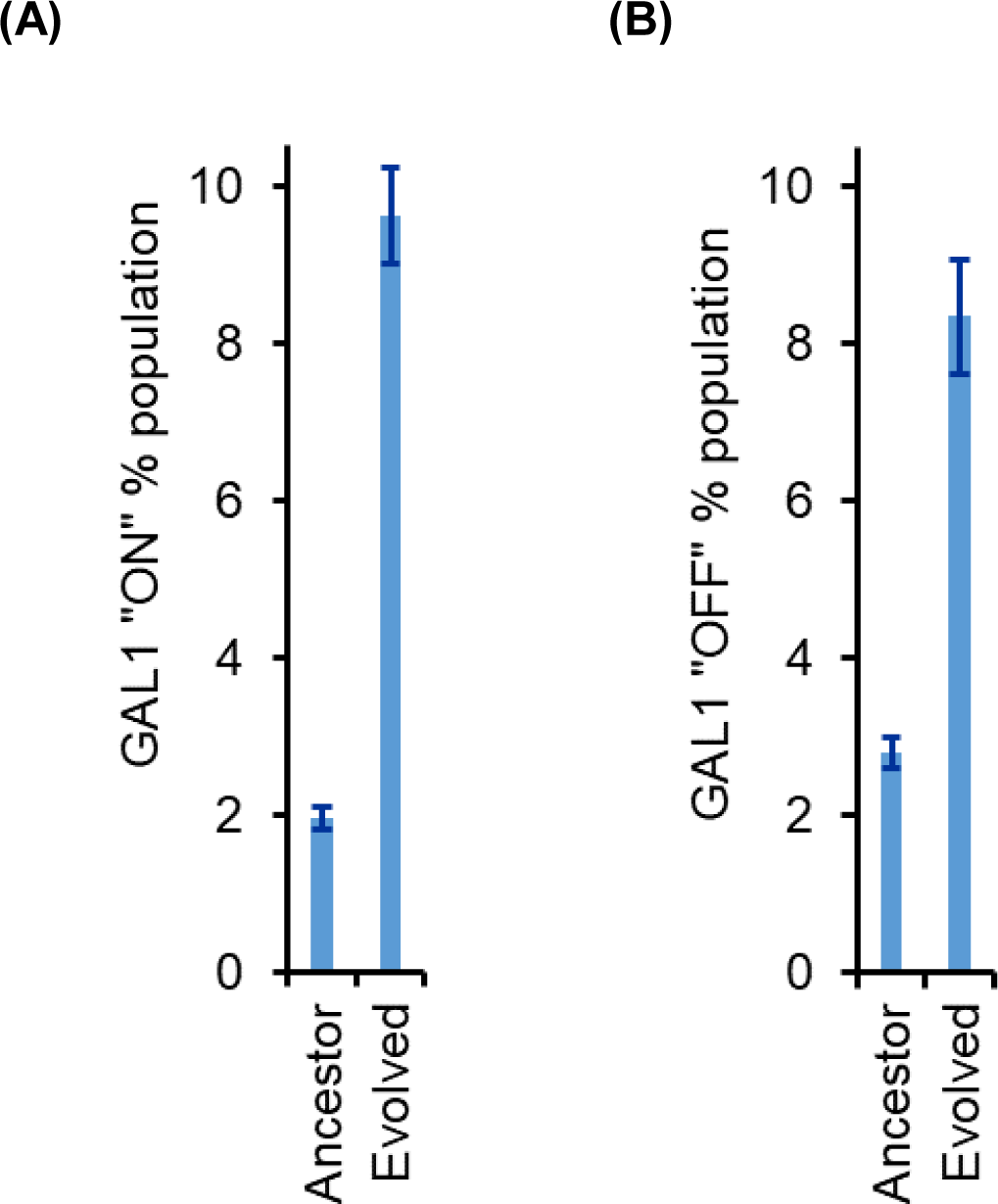
Percent population with low and high GAL1 induction increased in the evolved line (E1). **(A)** When mid-log phase culture growing on melibiose was spread on plates with xx concentration of 2DG, the evolved line (E1) had roughly 5-fold higher percent of population in GAL1 fully induced state. At this concentration of 2DG, only fully induced cells are killed. Percent population was calculated using the fraction of cells that did not survive on plates containing 2DG. **(B)** When mid-log phase culture growing on melibiose was spread on plates with xx concentration of 2DG, the evolved line (E1) had roughly 3-fold higher percent of population in GAL1 OFF state. This percent was calculated by counting the CFU on plates containing 2DG versus the CFU on YPD plates.

Thus, our results indicate that from the context of phenotypic metabolic state of the cells, both, ‘glucose-specialists’ and ‘galactose-specialists’ percentage in the evolved lines has increased. However, at the concentration of 2DG at which we test this resolution only identifies <10% of the ancestral, and <20% of the evolved populations. This is because the majority of the population expresses intermediate amounts of Gal1p.

Like the ancestor, the evolved line also exhibits colony size heterogeneity on melibiose plates. To study the differences in the small and large colony phenotype in the evolved line, we pick a small and a large colony from the evolved culture plated on melibiose. As a control, we pick a small and large colony from the ancestral culture also. Thereafter, we propagate these four colonies through a non-inducing non-repressing media (gly/lac) for about 15 generations (two 1:100 sub-cultures). Propagating cells in non-inducing non-repressing conditions for this duration was shown to have removed any differences between the metabolic state of the cells in small and large colonies (**Figure 6**).

After this period, the four cultures were plated on melibiose containing plates again. In the ancestral plate, the colony size distribution was identical between the cultures from small and large colonies. However, the colony size distribution from the large and small colonies of the evolved strain were qualitatively different from each other (**Figure 12**). The large colonies exhibited a much greater fraction of large colonies, as compared to the smaller one. As shown in the Figure, the colony size distribution was unimodal for the large and bimodal for small colonies from the evolved line E1.

**Figure 12.**
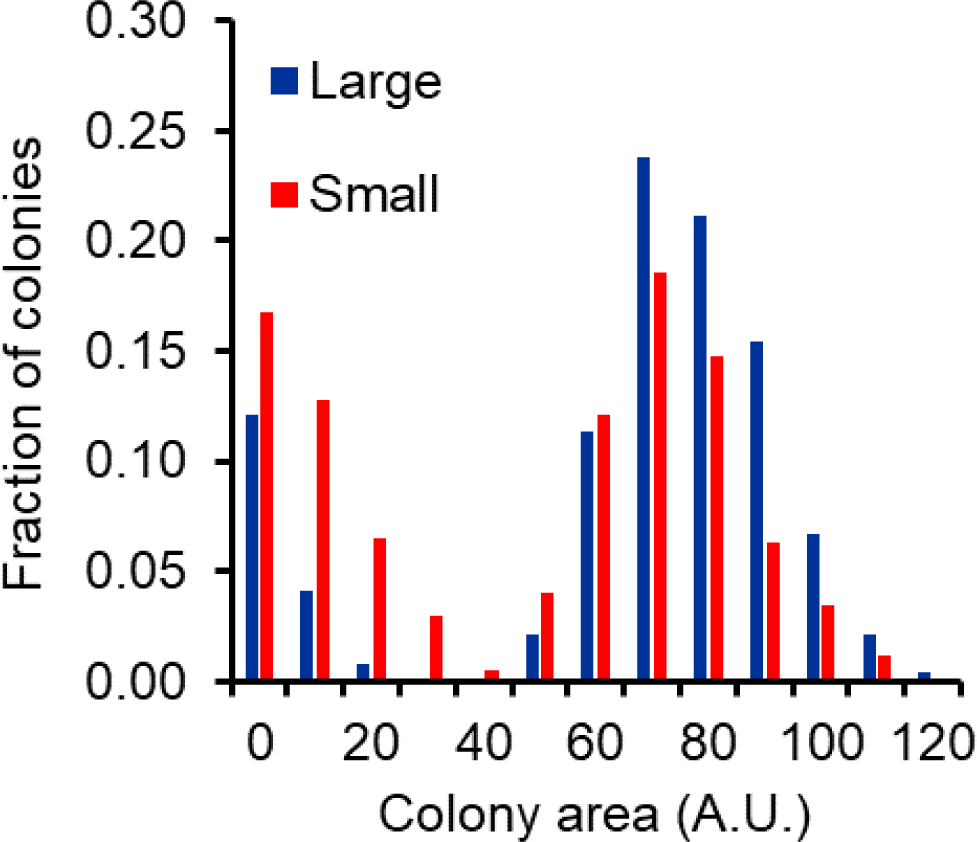
Large and small colonies in the evolved line E1 exhibit statistically significantly different colony size distribution on 1% melibiose plates. More than 1000 colonies were characterized for size for large and small colonies, when plated on melibiose.

Moreover, from the glycerol-lactate media, when the four colonies were tested for their growth on melibiose, the two cultures from the ancestral large and small colonies exhibit a growth dynamics identical to that of the ancestor. However, the large colony from the evolved culture exhibits a faster growth dynamics as compared to the small colony from the same evolved culture (**Figure 13**). The larger colony also exhibits faster growth in galactose, while the two exhibit similar growth kinetics in glucose. These results strongly suggest that during the course of the evolution experiment, the original isogenic population split into two groups, a glucose- and a galactose-specialist.

**Figure 13.**
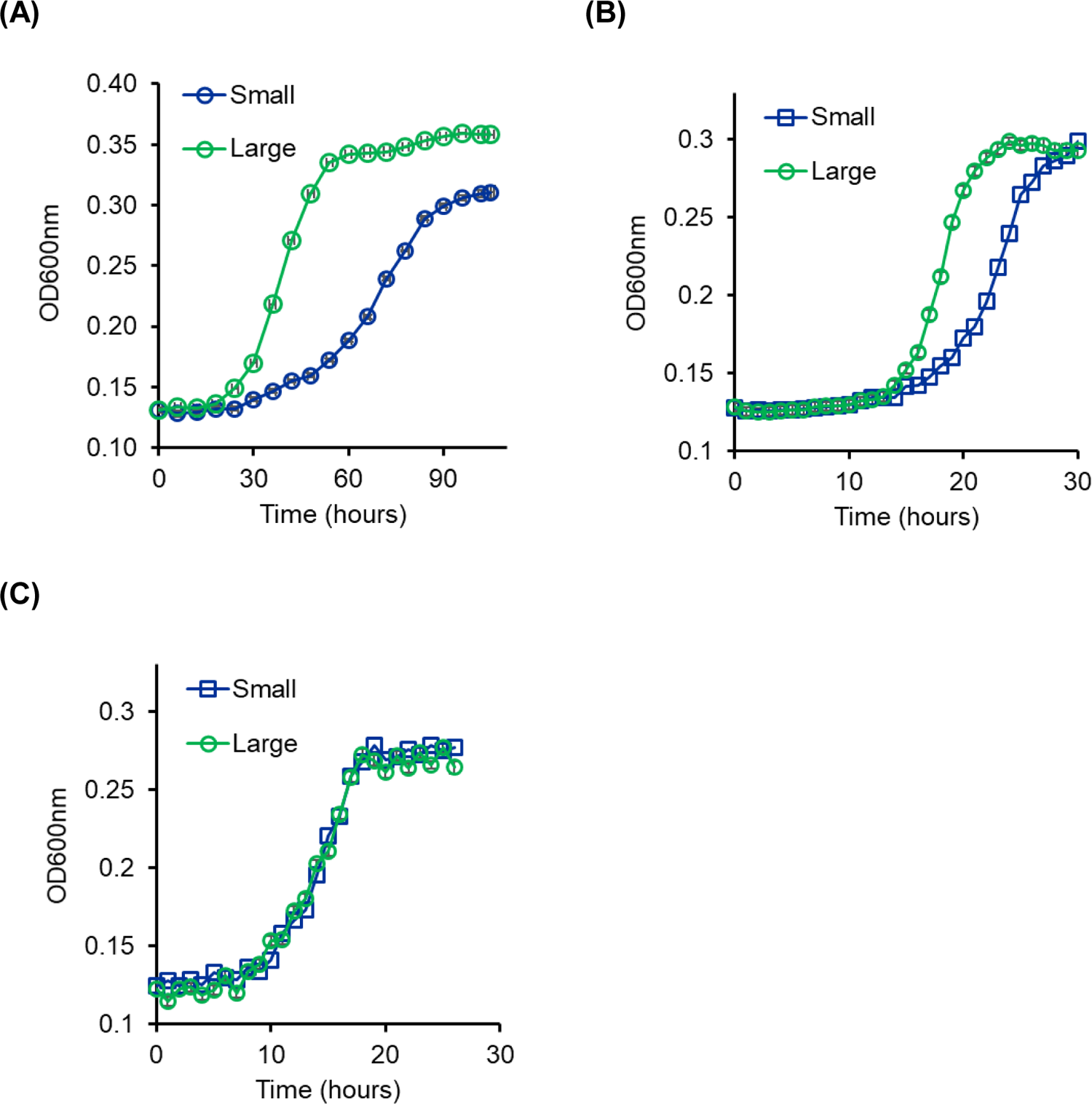
Evolved line E1 when plated on melibiose exhibits small and large colonies. The colonies exhibit different growth kinetics in 1% melibiose **(A)** and 1% galactose **(B)**. The colonies exhibit similar growth kinetics in 1% glucose **(C)**. To perform this experiment, small and large colonies from a melibiose plate were propagated in non-inducing non-repressing gly/lac media for 15 generations. The cultures were then diluted to an initial OD of 0.005 in the respective media and the growth kinetics monitored. The experiment was performed three times independently. The average and standard deviation are reported.

### Sequencing reveals mutation in the GAL3 locus in the large colonies on melibiose colonies

The coding and the promoter regions of *GAL1*, *GAL3*, *GAL4*, *GAL80*, and *MEL1* were sequenced. For this purpose, the evolved diploids (small and large colonies) were sporulated and the four haploids were isolated and the genes and the promoter regions requested. Sequencing results revealed that two mutations in the *GAL3* locus in the haploids isolated from large colony from the evolved strain. Previous work has demonstrated that difference in glucose-galactose signalling was attributed to the alleles present at the GAL3 locus [57]. As a result, the authors propose that GAL3 is a locus of high interest, as far as evolvability of the population is concerned in glucose-galactose mixtures.

Sequencing results show that in the large colony from the line E1, the *GAL3* locus had two mutations as compared to the ancestral sequence. These mutations (T363C and C1054G) lead to one synonymous and one non-synonymous changes in the coding sequence of *GAL3* (**Figure 14A**). Interestingly a recent analysis of *GAL3* sequence in environmental isolates of yeast *S. cerevisiae* revealed that these two mutations are also present in the strain NC-02 [57]. This particular strain carried two additional mutations in the coding region of *GAL3*. Moreover, the glucose repression on *GAL* gene expression in this particular strain was reported to be significantly lower as compared to that in the ancestral sequence (same as the ancestral sequence used in our study).

**Figure 14.**
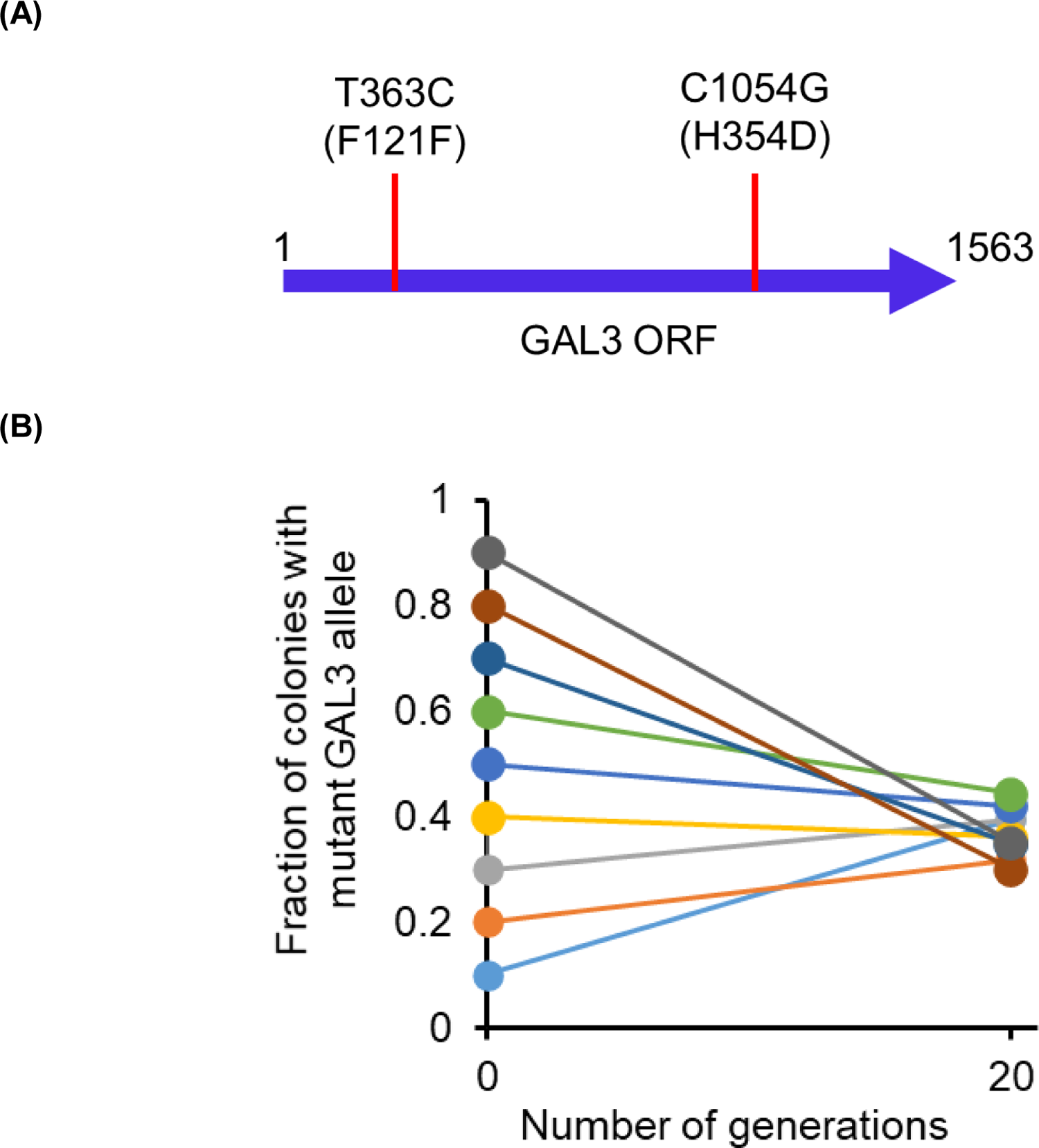
**(A)** Two SNPs in the coding sequence of the large colonies in the line E1. L121F was because of a T363C mutation, and H354D was because of a C1054G mutation. The second mutation (C1054G) is found in several yeast isolates. The mutation T363C is present in isolate NC-02 as described in reference [57]. **(B)** The evolved diploids (one carrying two ancestral *GAL3* allele and one carrying a mutant *GAL3* allele) were mixed in different ratios, and propagated in 1% melibiose for ∼20 generations (three 1:100 transfers). The relative ratio of the two strains converges to the unique value, independent of the starting point. The experiment was performed three independent times, and the average is represented. The error bar is smaller than the size of the data markers.

The mutant *GAL3* allele co-exists stably in a melibiose population along with the ancestor allele of *GAL3*. The evolved line when plated on melibiose plates, exhibits large and small colonies. The large colonies are from cells carrying the mutant copy of the *GAL3* allele. While the small colonies carry the ancestral *GAL3* allele. To check for co-existence, the small colonies (i.e., the evolved line carrying the ancestral *GAL3* allele) was transformed with a YCplac33 plasmid carrying an *URA3* allele. The two genotypes (evolved diploid with ancestor *GAL3* alleles carrying YCplac33, and the evolved diploid carrying a mutant *GAL3* allele) were mixed in different starting ratios and allowed to grow on melibiose for three sub-cultures (∼20 generations).

At the end of the growth period, an equal volume the cultures were plated on YPD plates and on ura^-^ synthetic media plates. The number of colonies on the two plates were used to estimate the relative ratio of the two alleles in the population. As shown in **Figure 14B**, after propagation in melibiose for ∼20 generations, the two strains converge to the same ratio. This ratio was found to be the same when the plasmid YCplac33 was transformed in the evolved diploid carrying the ancestral GAL3 allele (data not shown). Hence, the relative frequencies of the two evolved diploids is not because of the fitness load of the plasmid. And the two genotypes coexist in the melibiose media.

Gal3p protein is more than 70% identical with Gal1p, however, it lacks the galactokinase activity of Gal1p [58]. Gal3p has a phosphate-binding loop (spanning residues 156–162). This loop serves as the binding site for the ATP phosphoryl tail. The lack of galactokinase activity of Gal3p has been attributed to the absence of two amino acids in the GLSSSA(A/S)(F/L/I) motif typical of all functional galactokinases. Instead, Gal3p contains the sequence GLSSAF in that position. Gal3p can, however, be converted into a galactokinase through the addition of two amino acids, serine and alanine, after the Ser 164 residue in its coding region [59]. The two SNPs in the Gal3p coding region encode for a synonymous and non-synonymous change. Neither of the two SNPs impacts the region of the protein which is close to the active site. However, *GAL3* mRNA is known to be degraded faster when glucose is present in the media [60].

### Why not evolve to become better at utilizing melibiose?

Growth on melibiose is characterized by a relatively long lag phase, when compared to growth rate on monosaccharides glucose or galactose. The delay in growth, when *S. cerevisiae* is growing on melibiose, is due to the slow release of the enzyme α-galactosidase. The release of the enzyme in the media is considerably faster and higher in the evolved strain, as compared to the ancestor (**Figure 15**). It is known that promoters evolve fairly rapidly [61], and in this context, it is surprising that the *MEL1* promoter is weakly induced and has little constitutive activity. This is specially so since the fitness gain, in terms of reduced lag time, by increasing the *MEL1* promoter strength is significant. Why then is *MEL1* promoter so weakly induced? In our work, we do not observe any mutation in the strain where the *MEL1* promoter region has acquired any mutation.

**Figure 15.**
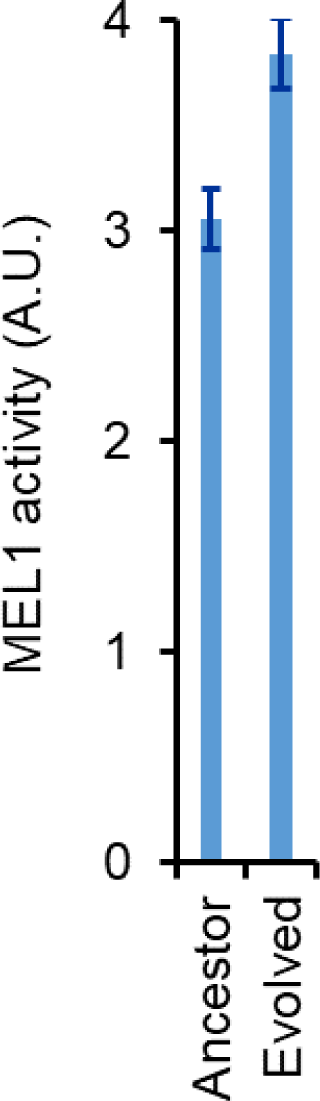
Evolved strain exhibits a higher Mel1p activity in the supernatant as compared to the ancestor.

We hypothesized that a stronger and a faster induction of the α-galactosidase from the *MEL1* promoter makes the strain more susceptible to cheater cells. To test this possibility, we performed competition experiments between the ancestor strain and BY4743. The *mel1*Δ strain is more than 40% fitter as compared to the ancestor strain, which carries a functional copy of *MEL1.* However, the strain BY4743 was more than 60% higher in fitness as compared to the ancestor. This demonstrates that improving the production and release of the α-galactosidase makes the population of *S. cerevisiae* to be more susceptible to exploitation by cheaters and other competitor strains. This disadvantage is likely important in an ecological niche (**Figure 16**).

**Figure 16.**
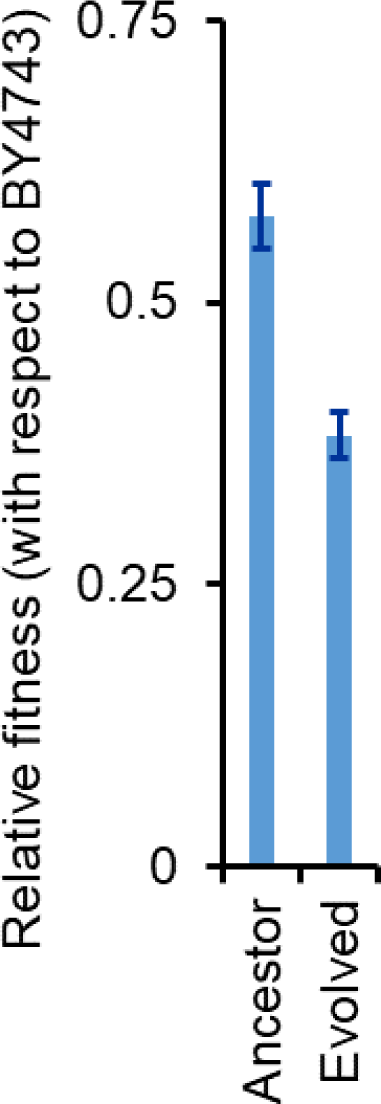
Fitness of the evolved line E1 is lower as compared to that of the ancestor. Competition experiments were performed between BY4743 (a mel1-strain) and the ancestor; and between BY4743 and the evolved line E1 in 1% melibiose. Relative fitness was calculated from the comparison of the relative frequency of the two competing strains at the start and end of the competition experiment.

A similar phenomenon, where production of a public good is decreased in order to avoid exploitation by cheaters has been observed previously. In Pseudomonas, when siderophore producers were evolved in the presence of cheater cells, in environments which contained low amounts of iron, the adaptive response of the siderophore producers (co-operators) was to lower the production of siderophores [62]. The fitness of these evolved co-operators was lower than that of the ancestor, when grown by themselves. However, in the presence of cheaters, the evolved strains were considerably fitter than the ancestor. Thus, reducing the public good production is an effective strategy in competing against cheater cells.

## Discussion

Isogenic cells exhibit a large variability in metabolic manifestations in steady state conditions. This difference has been attributed to gene expression noise [63]. Darwin proposed that diversification is achieved as specialists emerge to occupy all available niches, big or small, in a given environment [64]. Although, several examples of a population splitting into two distinct genotypes, and stably coexisting thereafter exist [5, 6, 65, 66] little is known about the fundamental of how this diversification happens [67].

Such a difference has been observed before – LTEE, and was noted in terms of colony size [68]. The underlying cause of divergence is availability of two resources, where specialization on one leads to reduced fitness on the other. However, in this example, the two resources are made available to the population in a temporal fashion, and are not simultaneously available [9, 69]. Divergence is also known to occur due to spatial heterogeneity [65, 70].

Antaogonism between galactose and glucose is known to occur via glucose-dependent inactivation of Rgt1p repressor [71, 72]; and galactose-dependent activation of the Rgt1 function, via activation of a co-repressor Mth1 [73]. Addition of glucose is also known to degrade *GAL3* mRNA, leading to enhanced expression of cyclin Cln3p – a key protein of cell division initiator [74]. These trade-offs have been shown to be an essential feature towards driving specialization in sympatric asexual populations [75].

Another dynamic feature of glucose-galactose utilization is that, in addition to diauxy, the greatest variability in *GAL* gene induction is seen when glucose is being consumed slowly, as compared to rapidly [76]. In the context of melibiose, since glucose is being released continuously, the rates of glucose depletion in the media are likely quite small. In the context of glucose to galactose transition, cell memory has been shown to be important in dictating the dynamics of transition [77]. While transition to galactose from non-inducing non-repressing environments like glycerol/lactate is relatively rapid and uniform; transition to galactose from glucose is slow and variable between different cells [78–80].

Several molecular targets could be involved in the diversification. From the context of the galactose network, variation in *GAL3* allele only is able to explain 90% variation in *GAL* induction kinetics among different environmental isolates [57]. *GAL3* was shown to be a locus, which modulates the diauxic lag, a selectable trait in the appropriate environmental conditions. To test this possibility, complementation experiments of the mutant *GAL3* allele, under its native and a constitutive promoter are being performed. The mutations in the *GAL3* allele are not close to the catalytic domain of the protein (analysis performed in Wincoot 0.9.4.1, results not shown). Thus, other possibilities of regulation remain to be tested. Gal3p is known to be degraded faster when glucose is present in the cell. One of the mechanistic explanations of the mutant Gal3p imparting greater growth on glucose could be that its growth rate is not enhanced when exposed to glucose. This would imply a greater Gal3p concentration in the cell, even in the presence of glucose, resulting in greater expression and release of Mel1p. This would lead to greater amounts of galactose and glucose being made available to the population, and hence, result in greater rates of growth, in both, galactose and melibiose.

### Author contributions

AM – evolution experiments, growth kinetic experiments,

2DG assay, competition experiments, α-galactosidase activity assay, wrote the manuscript.

PA – evolution experiments. VK – HPLC experiments.

RE – simulations.

PJB – conceived the study, analysed data.

SS – conceived the study, analysed data, wrote the manuscript.

## Acknowledgements

AM is supported by the Council of Scientific and Industrial Research (CSIR), Government of India, as a Senior Research Fellow (09/087(0873)/2017-EMR-I). The authors thank Barsa K. Godsora for help with analysis of the structure of the mutant GAL3 on Wincoot 0.9.4.1. The work was funded by DBT/Wellcome Trust India Alliance Grant IA/S/19/2/504632 to Supreet Saini.

